# Molecular Basis of PAC1R Allosteric Modulation with Lipids in Membranes

**DOI:** 10.1101/2025.01.27.635027

**Authors:** Nicholas B Hamilton, Bo Yang, Chijian Xiang, Tongtong Li, Severin T. Schneebeli, Jianing Li

## Abstract

The pituitary adenylate cyclase-activating polypeptide receptor I (PAC1R) represents a highly sought-after therapeutic target for chronic pain, migraine, and post-traumatic stress. As a class B G protein-coupled receptor (GPCR), the PAC1R is highly expressed throughout the neuronal and central nervous system membranes, with the receptor subject to hormone activation and subsequent signal transduction. Despite its desirable indication, small molecule agonists for PAC1R have been notoriously difficult to develop due to competition with PACAP. For this reason, allosteric activation of the receptor has emerged as a promising pathway. To probe potential allosteric sites, herein, we present a study of the receptor in biomimetic concentrations of lipid membranes. Our results reveal that cholesterol recognizes two canonical and two non-canonical binding sites at PAC1R, which may influence critical residues in the transmembrane domain and PAC1R activation. Also, our simulations suggest the glycolipid GM3 interacts with PAC1R in both the extracellular and transmembrane domains. These lipid binding hotspots may hold high potential for advancing our understanding of class B GPCR signaling and the discovery of new molecules targeting PAC1R.

**Statement of Significance:** The PAC1R is a highly sought after therapeutic target in the GPCR family. Understanding its natural process signaling is highly interesting in identifying new modes of control and is currently not well established. This work uses coarse-grained molecular dynamics simulations to examine diverse PAC1R models’ interactions with their endogenous lipid bilayer in compositions that match the regions in the brain where the receptor is expressed. Two lipids, cholesterol and GM3, have been previously identified in similar receptors as allosteric modulators and were specifically examined in this study. This work also showcases multiple new lipid binding sites at transmembrane and extracellular sites highly implicated in PAC1R signaling.

## Introduction

Secretin-like G-protein coupled receptors (also known as class B GPCRs) govern many key biological processes and represent potential targets (1, 2) for a wide class of physical and neurological disorders. These membrane receptor proteins, following the general taxonomy of all GPCRs, consist of seven helices forming a transmembrane domain (TMD), flanked on either side by chains and loops that shuttle signals and stimuli across the bilayer. Class B GPCRs have proven valuable targets for conditions such as type 2 diabetes (3), osteoporosis (4), and depression/post-traumatic stress disorder (5). Currently, several agonists (6, 7) and antagonists (8) of class B GPCRs are available on the market today. Thus, the mechanistic insight of class B GPCRs on the molecular level is fundamental for drug discovery and development.

The pituitary adenylate cyclase-activating polypeptide (PACAP) receptor (*ADCYAP1R1*, hereafter referred to as PAC1R) is a class B GPCR mainly expressed in the central and peripheral nervous system (9-11). PAC1R is activated by the endogenous PACAP peptide hormones to modulate the G-protein-dependent and independent pathways, involving transducers like G_αs_, G_αq_, and β-arrestin (12). The PACAP-bound structures of PAC1R in complex with heterotrimeric G_αs_ protein have been revealed by cryogenic electron microscopy (cryo-EM) (9). With multiscale simulations (13, 14), we have previously studied the conformational changes of ligand-free and β-arrestin-bound PAC1R (15, 16). In addition to the bent transmembrane helix 6 (TM6) in the active PAC1R conformation, the orientation and position of its extracellular domain (ECD) may also distinguish the active and inactive conformational states of PAC1R (15, 17). Further, our prior research (18) revealed that the *hop* isoform of PAC1R with a 28-amino acid *hop* insert in the intracellular loop 3 (ICL3), shows different signaling dynamics compared with the *null* isoform without any ICL3 insert (Figure S1). However, while GPCR-lipid interactions have been extensively investigated with class A receptors (19, 20), few class B receptors (21) have been studied, and there is still a knowledge gap in how isoforms in the active and inactive states are influenced by the lipid membrane environment.

Unlike ligand binding at the orthosteric pocket of a GPCR, the lipid contact is highly dynamic in nature, with both lipid re-organization and local GPCR side chain and backbone shifts in the conformational states. As such, the relative composition of these bilayers and the properties of each lipid component are of great interest to understanding GPCR conformational selection and the related signaling. In particular, such knowledge may lead to the discovery of new binding sites for allosteric modulators (22). Allosteric modulation of GPCRs can be a useful strategy for therapeutic development (23), with noted greater receptor selectivity and tunability for signaling control, compared to traditional GPCR small molecules targeting the orthosteric site (24, 25). Thus, it is helpful to use molecular dynamics (MD) simulations to examine the conformational transitions of GPCRs in the membrane (15), the detailed GPCR-lipid interactions in various membrane models (21), and GPCR interactions with signal transducers (26, 27). All-atom (AA) MD simulations can provide accurate details like hydrogen binding and charged interactions, while coarse-grained (CG) MD simulations can efficiently study and compare lipid-binding sites across multiple GPCR systems on longer timescales (28, 29). Combining AA and CG MD simulations, multiscale modeling (14, 30-32) can provide new insight into the structural information and dynamic nature of lipid binding. In this work, we focus on understanding the PAC1R interactions with cholesterol (CHOL) and monosialodihexosylganglioside (GM3) in the bilayer models and identifying different lipid preferences at active and inactive states of the GPCR. Cholesterol has been identified as an allosteric modulator in a host of transmembrane neuroreceptors (33, 34), exhibiting control over ligands that bind at the orthosteric site. A summary of GPCR structures co-crystalized with cholesterol in native recruitment sites is provided in Table S3, but as of yet, no sites for recruitment have been identified in the PAC1R. Highly abundant in the nervous system, glycolipids like GM3 have polar headgroups that extend above the membrane surface, which may interact with the extracellular loops or domains of GPCRs and stabilize specific confirmations (21, 28).

In this work, we have explored the PAC1R membrane dynamics with four GPCR constructs in two membrane models of up to eight different types of lipid-protein systems. The lipid concentrations were chosen to mimic brain tissues in areas where PAC1R is highly expressed (35, 36). Using CG MD simulations, we examined four constructs of PAC1 *hop* active, *hop* inactive, *null* active, and *null* inactive (Figure 1) to probe the connection between bilayer composition and the receptor-membrane dynamics of two key lipids indicated in allosteric modulation of GPCRs: cholesterol and GM3. Multiple key hydrophobic extrahelical recruitment sites were identified within the PAC1R for cholesterol, a known allosteric modulator of GPCRs (37). Interestingly, these sites at PAC1R share homology with previously reported class A GPCR complexes. In addition, the glycolipid GM3 was shown to interact with multiple sites within PAC1R, across the linker and extracellular domain depending on the activity of the receptor. We believe these sites hold high potential for advancing our understanding of class B GPCR signaling and the discovery of new molecules to modulate PAC1R for treating stress-related disorders (11).

**Figure 1.**
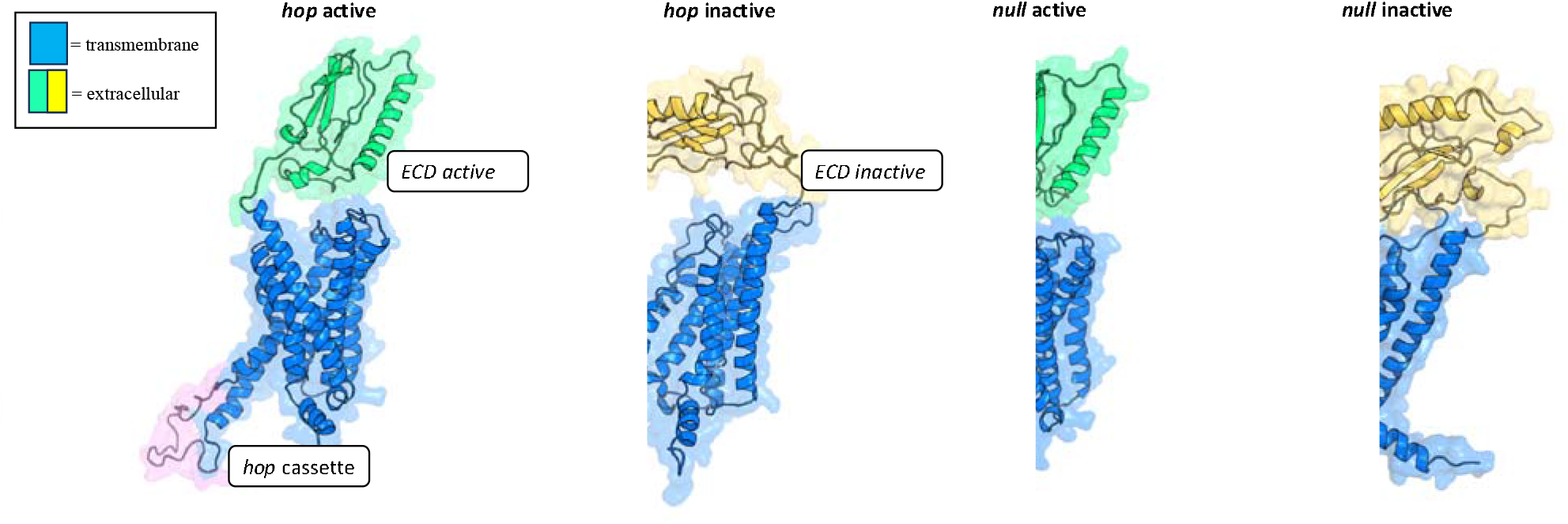
Cartoon illustration of four PAC1R models, featuring the transmembrane bundle of each isoform (blue), active ECD (green), inactive ECD (yellow), as well as the ICL3 hop insert (pink). Grey spheres illustrate the positions of the P atoms in the lipid bilayer.

## Methods and Models

### Model Construction

All MARTINI simulation inputs were prepared with the CHARMM-GUI MARTINI maker (38). Active and inactive models of the PAC1R *hop* and *null* isoforms were previously created using homology modeling (15), with the presence or absence of the 28-amino acid *hop* cassette to ICL3 (Figure 1). The presence of the *hop* cassette has previously been implicated in receptor activation within the brain specifically (39) and signaling dynamics of PAC1R (18). These four models (Figure 1) were embedded into the lipid bilayer membrane models, with the position suggested by the Orientation of Protein in Membranes (OPM) server (40). Two membrane models were considered (% by weight, Table S1-2): one symmetric bilayer model contains 21% cholesterol, 32% POPE, 29% POPC, 6% POPS, 5% BSM, 4% DPCE, and 3% POPI in both leaflets; the other model contains similar lipids in addition to 10% ganglioside GM3 in the upper leaflet. These compositions are based on the reported distribution of lipids within certain cross sections of the brain where PAC1R is highly expressed (35, 36). The MARTINI 2.0 force field (41) was used for the GPCR models and lipids, with the non-polarizable water model. The polar ElNeDyn (29) network was applied as constraints on the protein models. To confirm the contacts seen in the coarse-grained simulations, analogous AA models of each system were prepared using CHARMM-GUI. These systems were built to the same specifications as the MARTINI models and included the same lipid concentrations, disulfides. The CHARMM36 forcefield (42) was used for proteins and lipids in our AA models with the TIP3P water model.

### Coarse-Grained (CG) Simulation Setup

CG MD simulations were carried out using the GROMACS package (43). Steepest descent minimization was performed with the Verlet cutoff for neighbor searching. All minimization runs utilized Berendsen semi-isotropic pressure coupling with temperature coupling using velocity rescaling with a stochastic term within GROMACS. Five rounds of equilibration followed with the simulation in which force constraints were applied to the bilayer head groups and slowly removed with each successive step. The production runs were then performed in an NPT ensemble (300 K) with a 20-fs timestep and utilized the Parinello-Rahman semi-isotropic pressure coupling paired with the same temperature coupling scheme as the previous runs (velocity-rescale). Each production simulation was carried out for 20 μs, resulting in a total sampling of 160 µs across eight constructs (two PAC1R isoforms in two conformational states in two membrane models, as shown in Table S3).

### All-Atom (AA) Simulation Setup

AA simulations were carried out to validate our CG simulation observations of GM3 binding to PAC1R. The AA models were subject to a 10-ns Brownian dynamics simulation followed by a relaxation cycle of 100-ns simulation in an NVT ensemble with small force constraints on the heavy atoms. After relaxation, a series of three NPT steps were run with the force constraints progressively weakened. In the NPgT ensemble, we ran each Replica Exchange Molecular Dynamics (REMD) simulation using GROMACS 2024.1 (four replicas at temperatures of 303, 313, 323, and 343 K for 100 ns), with sampling of 1.6 μs from our AA simulations (Table S3).

### Model Visualization and Data Analysis

Our data analysis includes conformational analysis and lipid distributions. AA and CG models were visualized in PyMOL 3.0 (44). In-house Python scripts were developed using libraries such as MDAnalysis (45), pandas, numpy, and scikit-learn. PyMOL scripts (.pml) were created to match the per-residue lipid contacts to a color map. A per-residue contact is defined when the center-of-mass (COM) of cholesterol/GM3 and the COM of a residue are within 8.0 Å. Per-residue contacts were measured, normalized, and color-mapped onto the AA models for visualization to further examine each cholesterol or GM3 binding site. Each site was also mapped against the orthosteric pocket to identify which was most likely implicated in allosteric control from proximity to the pocket.

## Results and Discussion

### Cholesterol recognizes the canonical and non-canonical binding sites at PAC1R

From the rich simulation data provided by our CG models, we first measured the cholesterol distribution around our different PAC1 models. Cholesterol has been known to bind GPCRs at the so-called canonical and non-canonical binding sites (37). Cholesterol recognizes the GPCR surface with mostly hydrophobic interactions and hydrogen bonding, likely determined by the GPCR conformation and surface residues. On the other hand, cholesterol binding is believed to allosterically affect the GPCR conformations and activation (33, 37). To identify cholesterol-binding site hot spots (CHS) and compare different PAC1R models, the xy-positions of the lipids were mapped across all frames onto top-down color maps of each model (Figure 2). These results revealed four hotspots of PAC1R that mainly recruit cholesterol, namely CHS1 to CHS4. Of the four models examined, only the inactive form of the *null* isoform of the PAC1R strongly exhibited all four cholesterol recruitment sites from CHS1 to CHS4. CHS1 and CHS2 appear across all four models, in local proximity to one another across TM1 and TM2. The CHS1/CHS2 combined surface is likely dynamic, with cholesterol molecules aligning along the axis of the helices (Figure 3). From each of the four hotspots, we focus on the detailed interactions between cholesterol and five main amino acids: valine (V), threonine (T), leucine (L), phenylalanine (F), and isoleucine (I). Specifically, non-conservative V159, V166, and V173 in TM1 and V194, L198, I201, and I205 in TM2 contributed significantly to cholesterol binding at CHS1 and CHS2 (Figure 3).

**Figure 2.**
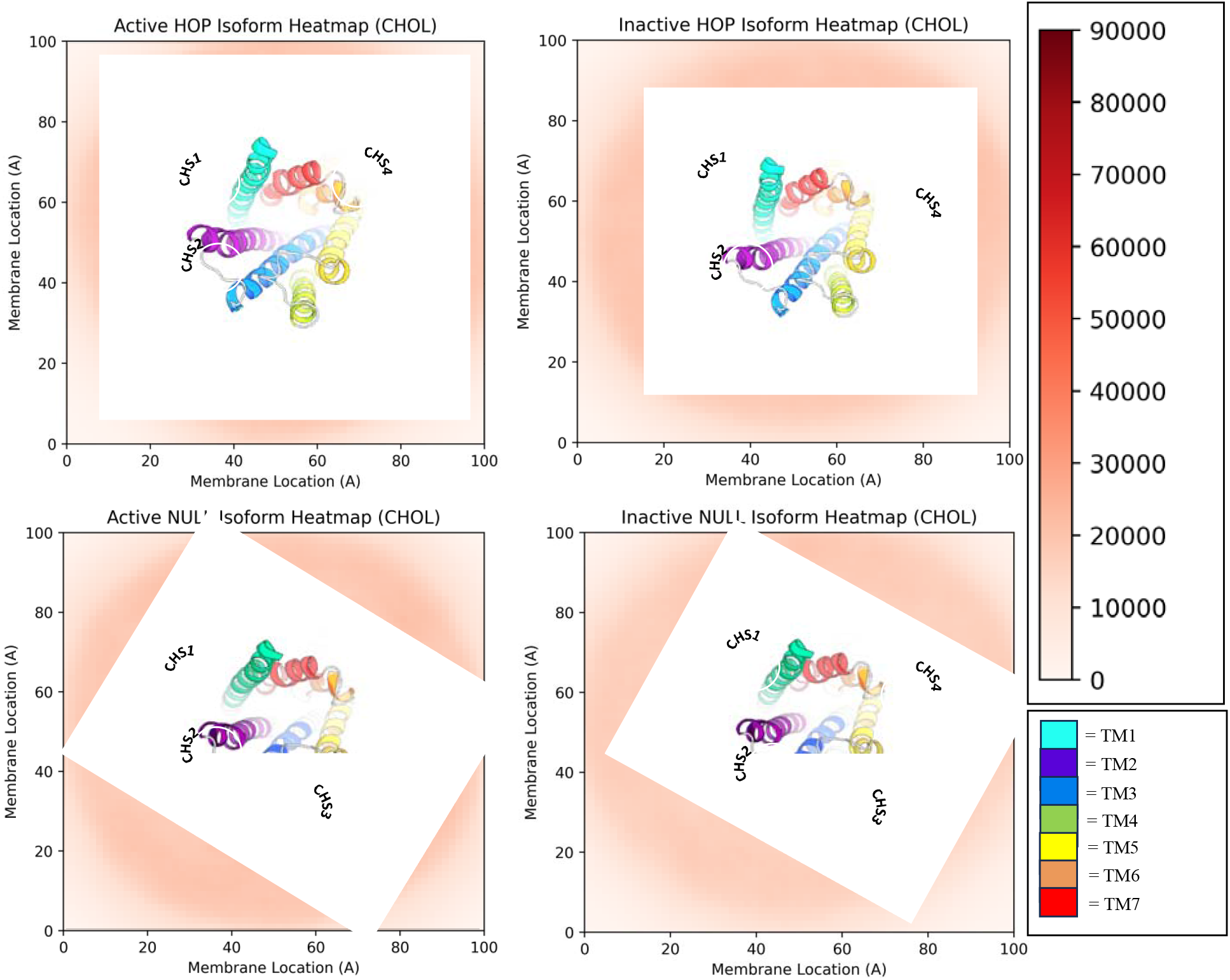
Top-down color maps of cholesterol distributions with the four PAC1R models (ECD not shown for clarity). The center of mass of each cholesterol molecule was mapped across the simulation. Four unique cholesterol-binding sites were recovered as the main correlation to the hotspot profile.

**Figure 3.**
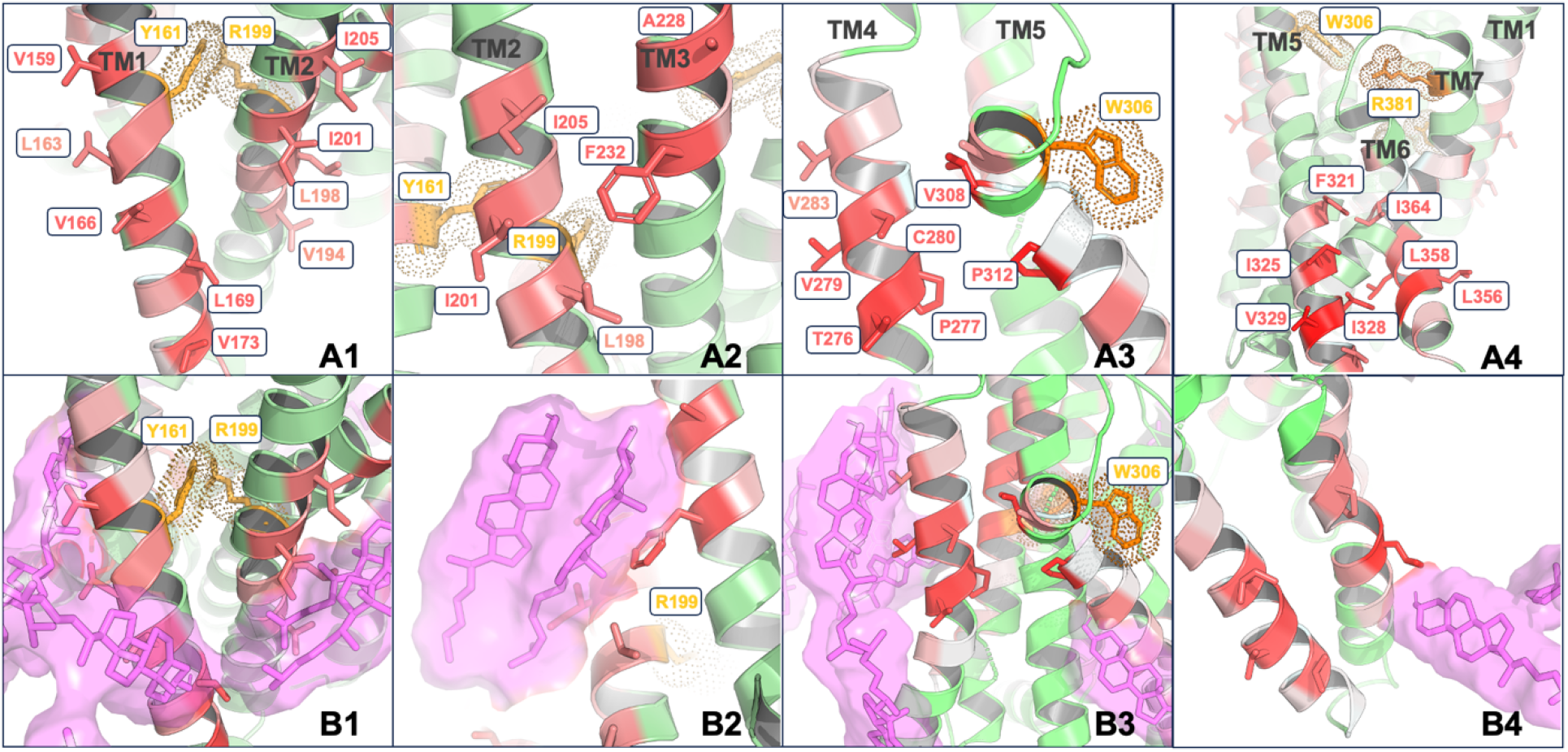
**(A1-4)** Color-coded active PAC1R models to illustrate the four cholesterol-binding hotspots from CHS1 to CHS4. PAC1R is represented in the green cartoon with cholesterol contacts scaling in red (deeper = more contacts). The four residues in the orthosteric pocket are highlighted in orange. **(B1-4)** Cholesterol (purple stick) positions backmapped to the all-atom model from the last frame of the corresponding CG simulation.

The PAC1R orthosteric site for PACAP binding was also examined regarding proximity to CHS1. This site has been extensively studied via hormone co-crystalized models (9, 46), revealing key residues for recruiting both the C and N termini of PACAP. From the PAC1R orthosteric pocket, we focused on four residues (Y161, R199, W306, R381) to measure proximity to the potential cholesterol hot spots. Thes four residues have previously been reported to control PACAP binding (11), with R381 and W306 rotated outward in the inactive model of PAC1R. The residues around CHS1 and CHS2 are shown to interweave two of these key orthosteric residues: Y161 and R199, which are known to play key roles in peptide binding and PAC1R activation (47). This observation indicates CHS1 and CHS2 as potential allosteric sites influencing PAC1R activation. Notably, these two sites also appeared to have high cholesterol densities across all four models, suggesting a dynamic binding motif as the receptor moves through signaling.

Along the extracellular side of TM4 and TM5, CHS3 appeared to be the only hot spot associated with the PAC1 *null* active and inactive models but suppressed in the corresponding *hop* variants (Figure 2). Like CHS1 and CHS2, CHS3 is near an active orthosteric site at W306, which may influence PAC1R activation allosterically. A further analysis identified V308 and P312 (Figure 3), which strongly recruited cholesterol without the *hop* insert. Furthermore, the intracellular site CHS4 appears as a cleft formed between TM5 and TM6 of PAC1R (Figure 3). Unlike the previous three sites, CHS4 is distant from the orthosteric site, existing on the intracellular side of the receptor. As ICL3 connects TM5 and TM6, the PAC1 *null* isoform with a shorter ICL3 likely displays higher sensitivity towards cholesterol binding to CHS3, while cholesterol shows stronger binding to the inactive PAC1R model at CHS4. A closer examination suggests that CHS4 has been observed in other GPCRs for cholesterol binding (33, 48, 49). For example, the CC motif chemokine 9 receptor has been shown to bind cholesterol at a site with analogous, pocket-like properties to CSH4 (PDBID: 5LWE) (50). In addition, CSH4 also contains substantial homology to the extrahelical binding site of another class B GPCR, the glucagon receptor (GCGR). MK-0893 (22) is a small-molecule antagonist bound to the extrahelical region of TM6 in GCGR (PDBID: 5EE7, Figure 4), which is adjacent to CHS4 observed in our simulations. This finding supports that the cholesterol hot spots can indicate potential allosteric binding sites, which may pave the way to develop new high-affinity allosteric modulators of PAC1R.

**Figure 4.**
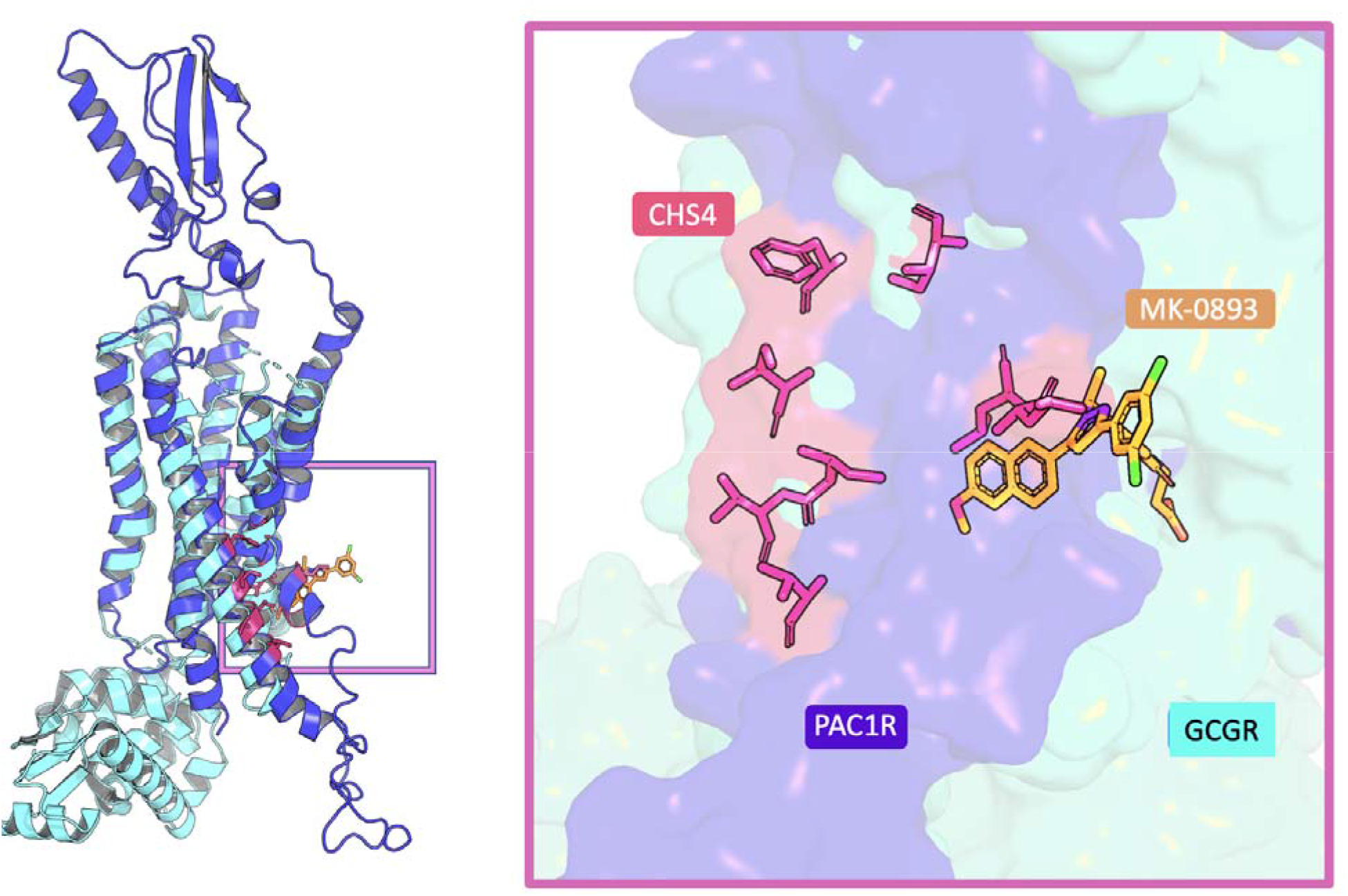
Left: The inactive PAC1 hop model (blue) overlapped with the inactive GCGR structure (cyan, PDBID 5EE7) bound to the negative allosteric modulator MK-0893 (orange stick). Right: Illustration of the non-canonical cholesterol binding site CHS4 on the PAC1R surface (blue), compared with the MK-0893 binding site on the GCGR surface (cyan).

The cholesterol recognition amino acid consensus (CRAC) (51) patterns like (K/R)-X_1–5_-(Y/F/W)-X_1–5_-(L/V) and (L/V)-X_1–5_-(Y/F/W)-X_1–5_-(K/R) were found in PAC1R. For example, CHS1 contains the CRAC patterns K154 -Y157-V159 and L192-F193-R199, while CHS2 has R199-F204-L210 and K227-F232-V237. Several crystal and cryo-EM structures show cholesterol bounds to CHS1 and CHS2 at the active and inactive GPCR structures (37). These two sites have been considered canonical for cholesterol binding. CHS3 and CHS4 appear affected by PAC1R isoforms or the activation states. CHS3 is near the outer leaflet of the membrane at TM5 and TM6, consistent with a non-canonical site in adenosine A2A receptor (PDBID: 4EIY and 5IU4) and several other class A GPCRs (Table S3). CHS4 is in the inner leaflet of the membrane at TM5, TM6, and TM7, close to the non-canonical site observed at TM1 and Helix 8 of 5-Hydroxytryptamine receptor 2B (PDBID: 4IB4, 5TVN, and 6DRX). In short, our simulation results show cholesterol recognizes the canonical (CHS1 and CHS2) and non-canonical (CHS3 and CHS4) binding sites at PAC1R.

### GM3 binding is affected by PAC1R ECD

We examined the local binding affinity of the glycolipid GM3, which has been implicated in modulating the progression of the GCGR along its conformational ensemble (21). GM3 has been shown to aggregate around both the linker and flexible ECD region of the GCGR, affecting the motion of the ECD and subsequent population of conformational states as it progresses through signaling (21). As such, the GM3 distribution is useful for examining the role of membrane-protein dynamics related to PAC1R. PAC1R may follow a complex activation pathway (15), which transitions quickly between several ECD closed states but has a long transition from any closed state to the ECD open state without PACAP. During these transitions, the linker is highly flexible. As a result, ECD can be positioned directly above the membrane and likely affected by GM3 recruitment at both the linker interface controlling the ECD orientation and the exposed surface residues within the ECD itself. In addition, GM3 has previously been reported to be recruited selectively by other GPCRs at sites homologous to the PAC1R ECD and the linker (21). To visualize the GM3 distribution near PAC1R, we mapped the xy-position of the center of mass of each lipid of interest and projected them onto the color maps (Figure 5). GM3 binding hotspots were identified as GHS1, GHS2, GHS3, and GHS4. Compared with cholesterol, GM3 interactions with PAC1R greatly depend on the ECD position and orientation.

**Figure 5.**
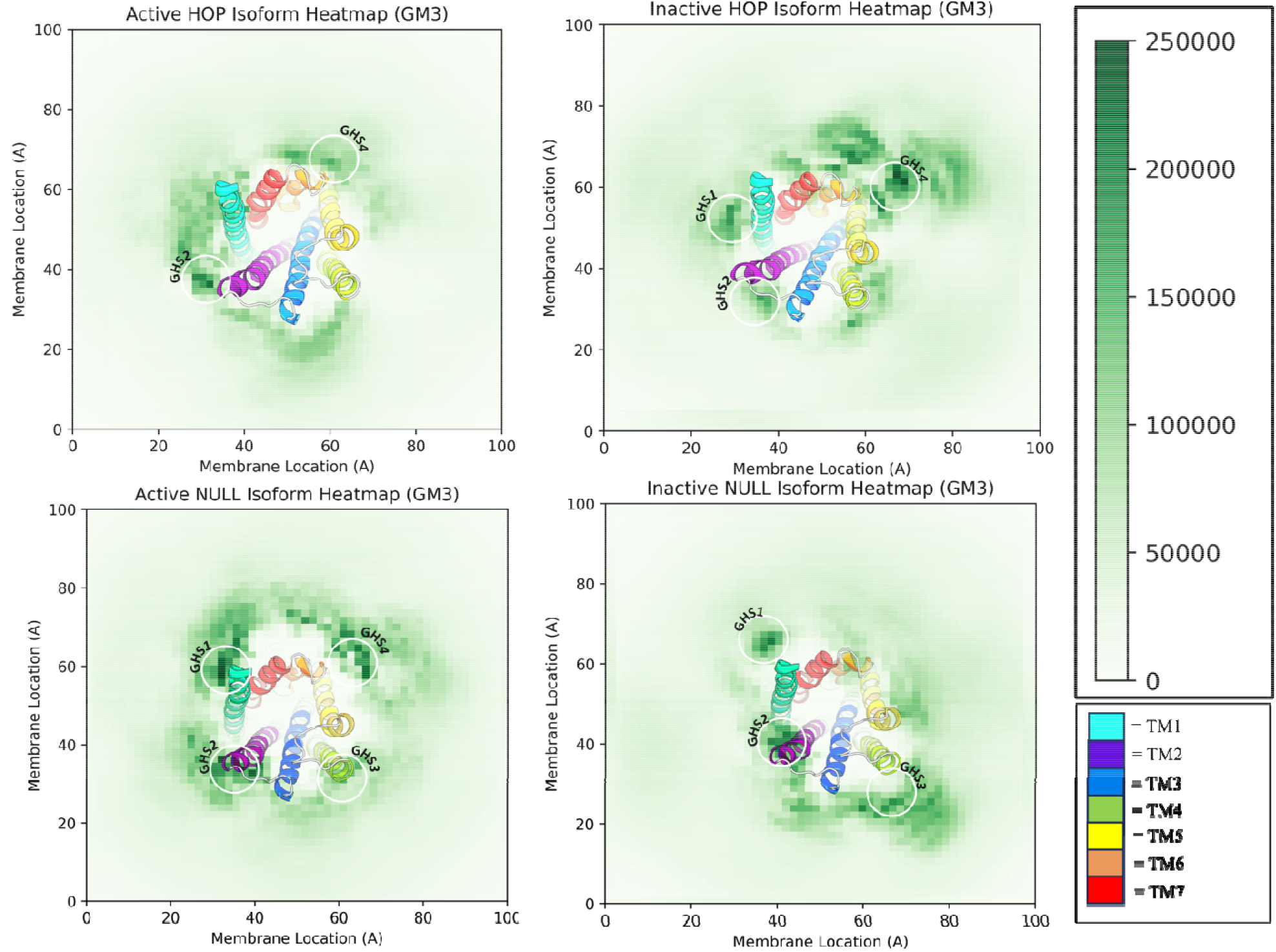
Top-down color maps of cholesterol distributions with the four PAC1R models (ECD not shown for clarity). The center of mass of each GM3 molecule was mapped across the simulation. Four unique GM3-binding sites were recovered as the main correlation to the hotspot profile.

In our coarse-grained models, GM3 is shown to exhibit diverse recruitment at new sites of the PAC1R dependent on the receptor conformations. The first of these hot spots, dubbed GHS1, was shown to appear across all four models as an extended combination of residues shared across the extracellular loops between TM2, TM3, and TM4. GHS1 appeared in the inactive PAC1R models across the ECL loops, with the addition of the kink in TM5 shown to provide high recruitment and extend the overall site compared to both the active models. GHS1 was the overall favored site for GM3 within the transmembrane domain across all four of our PAC1R models. Our simulations also revealed complex recruitment dynamics between the ECD/linker and GM3. In both inactive models, PAC1R is shown to heavily recruit GM3 at the linker site (namely GHS2), with residues above TM1 heavily implicated (Figure 6). This activity trend is reversed in the complimentary active PAC1R models, with the ECD and linker shown to recruit very little GM3 in comparison. Of the residue found to interact with GM3, E142 and T143 were shown to be key in the inactive models while recording no contacts in their complimentary active form. GHS2 confirms the selective recruitment of GM3 displayed in ECD and linkers of the inactive models of PAC1R.

**Figure 6.**
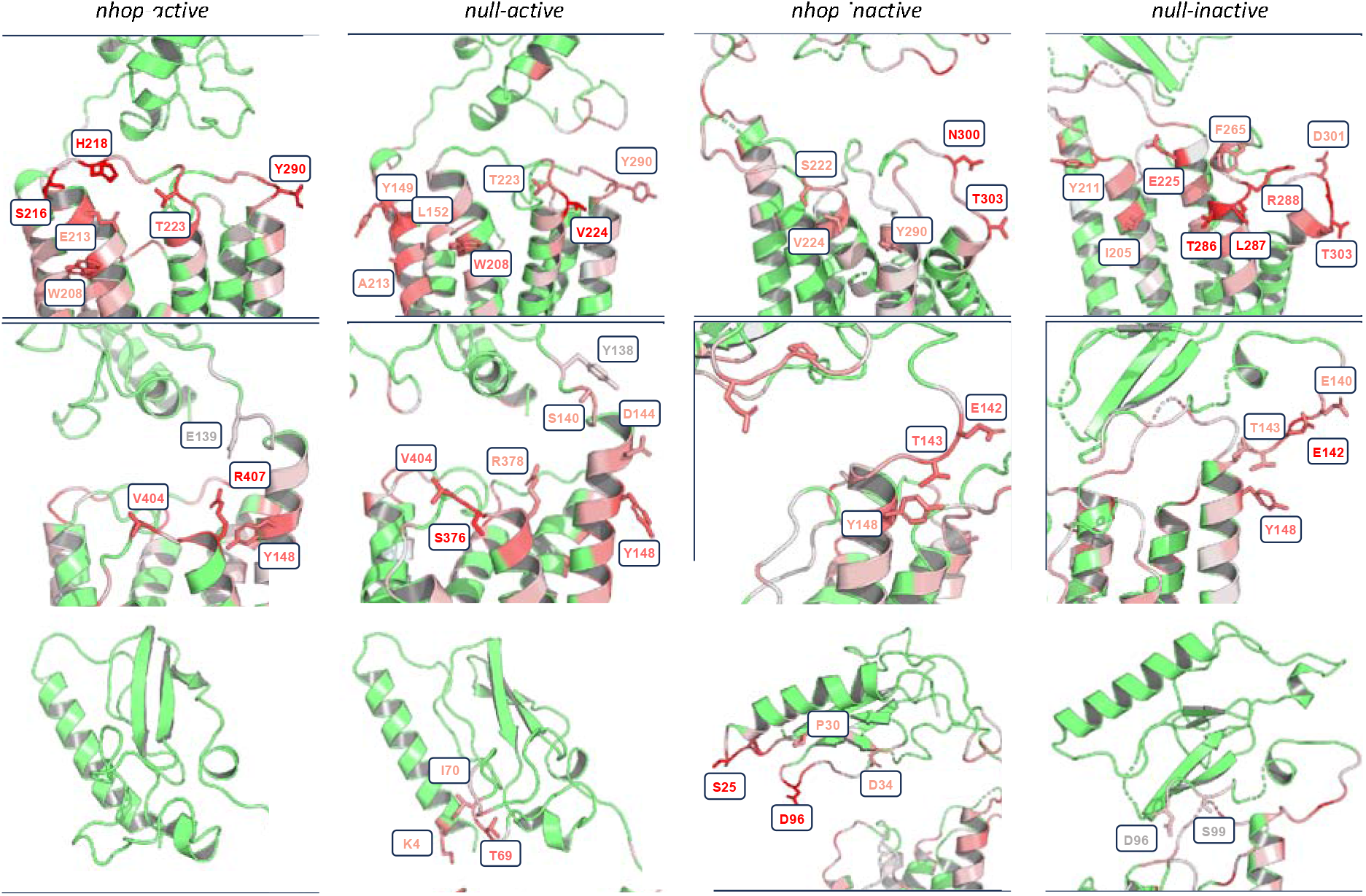
GM3 lipid binding hot spots mapped onto the PAC1R models. **(A)** GHS1 shows an extended motif along the extracellular loops of all four models, with the kink in the inactive forms shown to extend the recruitment site for the inactive models. **(B)** GHS2 at the interface of the extracellular loops of TM5/TM6 and the linker to the PAC1R ECD. **(C)** GHS3 within the PAC1R ECD. Recruitment is shown to be ECD orientation independent, with both active and inactive models shown to bind GM3.

In addition to the linker recruitment selectivity, we identified one more feature displayed by our models: the recruitment of GM3 in the ECD domain (Figure 6C). Of the models sampled, only the *null* active and *hop* inactive forms of the PAC1R were shown to recruit substantial amounts of GM3, with many surface residues shown to recruit the glycolipid at an equivalent rate to the TM site GHS1. Notably, the position of the ECD and subsequent exposure of the PAC1R N-terminus was heavily favored in the active model. This resulted in lysine K4 being the most exposed towards GM3 in the active model within GHS3. In the inactive state, S25 and D29 are the corresponding residues within the inactive ECD. These two were shown to recruit GM3 at an analogous solvent-exposed site to K4 in the corresponding active model (Figure 6C). The *null* inactive model also showcased minor recruitment of GM3 to its ECD throughout the solution, albeit at a much lower rate than the contacts in the active complementary model. Despite the profound absence of GM3 in the ECD of the *hop* active model, our data does not suggest that there is a strong correlation between receptor activity and recruitment at GHS3. The sequence of the models with the *hop* insert to ICL3 also did not appear correlated with this site nor GHS1 or GHS2. Considering the presence of the insert on ICL3, isolated from any of the GM3 hotspots, combined with the elastic constraints applied from the MARTINI CG model restricting the receptor’s structural evolution, further study is required to elicit the ECD effects from GM3 binding. As shown previously, cholesterol appeared affected by the cassette, with CHS3 selective for the *null* isoform over the *hop*. We believe cholesterol to be a more transient partner in lipid-PAC1R affinity and, thus, can access a wider variety of states and sites based on topology. GM3, on the other hand, we conclude to be heavily linked to the conformation state of the ECD within PAC1R. As such, GM3 hotspots were seen to be more affected by activity, with the inactive models specifically sharing almost identical recruitment at sites GHS1 and GHS2 between isoforms. Internalization of GM3 within the ECD itself is possible for both the active and inactive state of PAC1R, and corresponding solvent-exposed residues exist within each model for GM3 recruitment.

While the results of our coarse-grained simulations show the link between membrane composition and receptor dynamics, we further sought to validate that these new contacts were not spawned from bias in our initial models and maintained by the CG constraints provided for receptor stability by the ElNeDyn network. We performed REMD simulations of the AA models assembled to the same concentrations as the CG simulations to validate our findings. We calculated contacts with lipids of interest for each residue from these models and mapped the scaled values back onto a 3D model for comparison with the CG simulation results (Figure S2). The active and inactive forms of the *null* isoform were shown to confirm GM3 recruitment in the ECD (GHS3) as a motif for the inactive state. In addition, GHS1 and GHS2 were shown to appear in the REMD models, though of varying intensity and overlap due to the reduced simulation time.

## Conclusions

In conclusion, systematic CG simulations of the PAC1 receptor have revealed complex lipid-protein dynamics. Two functional lipids, cholesterol and GM3, were shown to have different interactions towards multiple local sites on the PAC1R surface. Cholesterol binds dynamically based on the receptor activity and sequence identity. Of the four spots where cholesterol was recruited within PAC1R, CHS1 appears to be the most promising for allosteric control, as it overlaps with R199 and Y161, which are key binding partners for PACAP, the receptor’s endogenous hormone. In addition, CHS4 was shown to recruit cholesterol and share homology with the small molecule binding site at a closely related class B GPCR. A potential mechanistic motif for advancing PAC1R’s ECD along its previously reported conformational pathway modulated by GM3 was also observed in the inactive state of both the *hop* and *null* isoforms of PAC1R. This motif was observed through the enhanced binding of GM3 to both the linker and ECD of the inactive PAC1R models compared to the complimentary active states. This result was confirmed with all-atom REMD of receptor models recreated in an all-atom environment to the same specification as the coarse-grained simulations. These AA models revealed a similar mechanism of ECD lipid recruitment as seen in the CG simulations and general overlap for the recruitment sites (Figure S4). GM3 will likely play a key role in furthering this pathway and subsequent PACAP binding, as this open-closed receptor transition has been linked. These findings can be useful to guide further design of allosteric modulators to selectively modulate PAC1R activity and provide new solutions for PAC1R-related conditions like stress and chronic pain.

## Acknowledgments

We thank Drs. Ray Bressan (Purdue) and Jacob Remington (UVM) for helpful discussions. The work was mainly supported by an NIH grant R01-GM129431 to J.L. N.B.H. was partially supported by an NSF award (CHE-1945394/2410514 to J.L.). S.T.S. was supported by an NIH R35 award (R35-GM147579).

