## Supplemental information for "Molecular Basis of PAC1R Allosteric Modulation with Lipids in Membranes"

Figure S2..…………………………………………………………………………………………………………………………….. 10

**Table S1. Composition of Lipid Membrane without glycolipids.**

| Lipid | Weight % |
| --- | --- |
| CHOL | 21 |
| DPCE | 4 |
| POPE | 32 |
| POPC | 29 |
| POPS | 6 |
| POPI | 3 |
| BSM | 5 |

**Table S2. Composition of Lipid Membrane with glycolipids**

| Lipid | Weight %  *Upper leaflet* | Weight %  *Lower leaflet* |
| --- | --- | --- |
| CHOL | 19 | 21 |
| DPCE | 3 | 4 |
| POPE | 30 | 32 |
| POPC | 28 | 29 |
| POPS | 5 | 6 |
| POPI | 3 | 3 |
| BSM | 2 | 5 |
| GM3 | 10 | 0 |

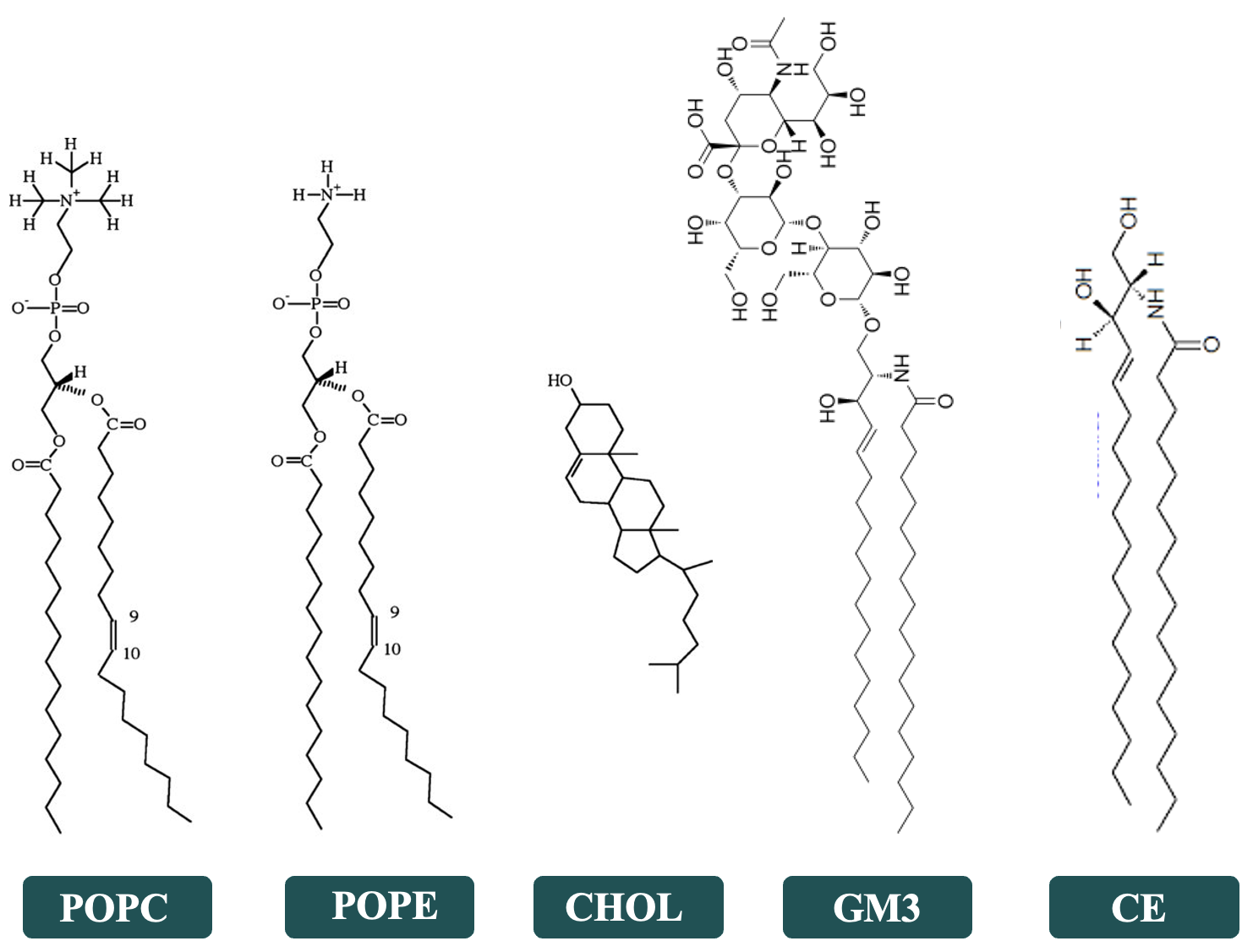

**Table S3. Simulation Summary. Table S1 shows the S1 membrane lipid composition; Table S2 shows the S2 membrane lipid composition.**

| PAC1R Construct | Membrane Models | Number of  CG sites/Atoms | Box Size (x, y, z) in Å | Simulation Length (μs) | Temperature (K) |
| --- | --- | --- | --- | --- | --- |
| CG, hop (active) | S1 | 13,226 | 100, 100, 150 | 20 | 303 |
| CG, hop (inactive) | S1 | 12,893 | 100, 100, 150 | 20 | 303 |
| CG, null (active) | S1 | 12,834 | 100, 100, 150 | 20 | 303 |
| CG, null (inactive) | S1 | 13,393 | 100, 100, 150 | 20 | 303 |
| CGhop (active) | S2 | 12,446 | 100, 100, 150 | 20 | 303 |
| CG, hop (inactive) | S2 | 13,885 | 100, 100, 150 | 20 | 303 |
| CG null (active) | S2 | 13,547 | 100, 100, 150 | 20 | 303 |
| CG, null (inactive) | S2 | 12,750 | 100, 100, 150 | 20 | 303 |
| AA, hop (active) | S2 | 155,687 | 100, 100, 150 | 0.1×4 | 303. 313, 323, 333 |
| AA, hop (inactive) | S2 | 153,681 | 100, 100, 150 | 0.1×4 | 303. 313, 323, 333 |
| AA, null (active) | S2 | 153,570 | 100, 100, 150 | 0.1×4 | 303. 313, 323, 333 |
| AA, null (inactive) | S2 | 144,073 | 100, 100, 150 | 0.1×4 | 303. 313, 323, 333 |

**Table S4. Summary of GPCRs-Cholesterol Complexes in PDB.**

| PDBID | Structure Title | Experiment |
| --- | --- | --- |
| 2RH1 | High resolution crystal structure of human B2-adrenergic G protein-coupled receptor. | X-RAY |
| 3D4S | Cholesterol bound form of human beta2 adrenergic receptor. | X-RAY |
| 3NY8 | Crystal structure of the human beta2 adrenergic receptor in complex with the inverse agonist ICI 118,551 | X-RAY |
| 3NY9 | Crystal structure of the human beta2 adrenergic receptor in complex with a novel inverse agonist | X-RAY |
| 3NYA | Crystal structure of the human beta2 adrenergic receptor in complex with the neutral antagonist alprenolol | X-RAY |
| 3PDS | Irreversible Agonist-Beta2 Adrenoceptor Complex | X-RAY |
| 4EIY | Crystal structure of the chimeric protein of A2aAR-BRIL in complex with ZM241385 at 1.8A resolution | X-RAY |
| 4NTJ | Structure of the human P2Y12 receptor in complex with an antithrombotic drug | X-RAY |
| 4PXZ | Crystal structure of P2Y12 receptor in complex with 2MeSADP | X-RAY |
| 5D5A | In meso in situ serial X-ray crystallography structure of the Beta2-adrenergic receptor at 100 K | X-RAY |
| 5D5B | In meso X-ray crystallography structure of the Beta2-adrenergic receptor at 100 K | X-RAY |
| 5IU4 | Crystal structure of stabilized A2A adenosine receptor A2AR-StaR2-bRIL in complex with ZM241385 at 1.7A resolution | X-RAY |
| 5IU7 | Crystal structure of stabilized A2A adenosine receptor A2AR-StaR2-bRIL in complex with compound 12c at 1.9A resolution | X-RAY |
| 5IU8 | Crystal structure of stabilized A2A adenosine receptor A2AR-StaR2-bRIL in complex with compound 12f at 2.0A resolution | X-RAY |
| 5IUA | Crystal structure of stabilized A2A adenosine receptor A2AR-StaR2-bRIL in complex with compound 12b at 2.2A resolution | X-RAY |
| 5IUB | Crystal structure of stabilized A2A adenosine receptor A2AR-StaR2-bRIL in complex with compound 12x at 2.1A resolution | X-RAY |
| 5L7D | Structure of human Smoothened in complex with cholesterol | X-RAY |
| 5D6L | beta2AR-T4L - CIM | X-RAY |
| 5K2A | 2.5 angstrom A2a adenosine receptor structure with sulfur SAD phasing using XFEL data | X-RAY |
| 5K2B | 2.5 angstrom A2a adenosine receptor structure with MR phasing using XFEL data | X-RAY |
| 5K2C | 1.9 angstrom A2a adenosine receptor structure with sulfur SAD phasing and phase extension using XFEL data | X-RAY |
| 5K2D | 1.9A angstrom A2a adenosine receptor structure with MR phasing using XFEL data | X-RAY |
| 5LWE | Crystal structure of the human CC chemokine receptor type 9 (CCR9) in complex with vercirnon | X-RAY |
| 5TVN | Crystal structure of the LSD-bound 5-HT2B receptor | X-RAY |
| 5JTB | Crystal structure of the chimeric protein of A2aAR-BRIL with bound iodide ions | X-RAY |
| 5MZJ | Crystal structure of stabilized A2A adenosine receptor A2AR-StaR2-bRIL in complex with theophylline at 2.0A resolution | X-RAY |
| 5MZP | Crystal structure of stabilized A2A adenosine receptor A2AR-StaR2-bRIL in complex with caffeine at 2.1A resolution | X-RAY |
| 5N2R | Crystal structure of stabilized A2A adenosine receptor A2AR-StaR2-bRIL in complex with PSB36 at 2.8A resolution | X-RAY |
| 5NLX | A2A Adenosine receptor room-temperature structure determined by serial millisecond crystallography | X-RAY |
| 6AQF | Crystal structure of A2AAR-BRIL in complex with the antagonist ZM241385 produced from Pichia pastoris | X-RAY |
| 5OLG | Structure of the A2A-StaR2-bRIL562-ZM241385 complex at 1.86A obtained from in meso soaking experiments. | X-RAY |
| 5OLH | Structure of the A2A-StaR2-bRIL562-Vipadenant complex at 2.6A obtained from in meso soaking experiments. | X-RAY |
| 5OLO | Structure of the A2A-StaR2-bRIL562-Tozadenant complex at 3.1A obtained from in meso soaking experiments. | X-RAY |
| 5OLV | Structure of the A2A-StaR2-bRIL562-LUAA47070 complex at 2.0A obtained from in meso soaking experiments. | X-RAY |
| 5OLZ | Structure of the A2A-StaR2-bRIL562-Compound 4e complex at 1.9A obtained from bespoke co-crystallisation experiments. | X-RAY |
| 5OM1 | Structure of the A2A-StaR2-bRIL562-Compound 4e complex at 2.1A obtained from in meso soaking experiments (1 hour soak). | X-RAY |
| 5OM4 | Structure of the A2A-StaR2-bRIL562-Compound 4e complex at 1.86A obtained from in meso soaking experiments (24 hour soak). | X-RAY |
| 6B73 | Crystal Structure of a nanobody-stabilized active state of the kappa-opioid receptor | X-RAY |
| 6D35 | Crystal structure of Xenopus Smoothened in complex with cholesterol | X-RAY |
| 5WB2 | US28 bound to engineered chemokine CX3CL1.35 and nanobodies | X-RAY |
| 6DRX | Structural Determinants of Activation and Biased Agonism at the 5-HT2B Receptor | X-RAY |
| 6DRY | Structural Determinants of Activation and Biased Agonism at the 5-HT2B Receptor | X-RAY |
| 6DRZ | Structural Determinants of Activation and Biased Agonism at the 5-HT2B Receptor | X-RAY |
| 6DS0 | Structural Determinants of Activation and Biased Agonism at the 5-HT2B Receptor | X-RAY |
| 6A93 | Crystal structure of 5-HT2AR in complex with risperidone | X-RAY |
| 6A94 | Crystal structure of 5-HT2AR in complex with zotepine | X-RAY |
| 6GT3 | Crystal Structure of the A2A-StaR2-bRIL562 in complex with AZD4635 at 2.0A resolution | X-RAY |
| 6IBB | Crystal structure of the rat isoform of the succinate receptor SUCNR1 (GPR91) in complex with a nanobody | X-RAY |
| 6JZH | Structure of human A2A adenosine receptor in complex with ZM241385 obtained from SFX experiments under atmospheric pressure | X-RAY |
| 6PRZ | XFEL beta2 AR structure by ligand exchange from Alprenolol to Alprenolol. | X-RAY |
| 6PS0 | XFEL beta2 AR structure by ligand exchange from Alprenolol to Carazolol. | X-RAY |
| 6PS1 | XFEL beta2 AR structure by ligand exchange from Alprenolol to Timolol. | X-RAY |
| 6PS2 | XFEL beta2 AR structure by ligand exchange from Timolol to Alprenolol. | X-RAY |
| 6PS3 | XFEL beta2 AR structure by ligand exchange from Timolol to Carvedilol. | X-RAY |
| 6PS4 | XFEL beta2 AR structure by ligand exchange from Timolol to ICI-118551. | X-RAY |
| 6PS5 | XFEL beta2 AR structure by ligand exchange from Timolol to Propranolol. | X-RAY |
| 6PS6 | XFEL beta2 AR structure by ligand exchange from Timolol to Timolol. | X-RAY |
| 6PS7 | XFEL A2aR structure by ligand exchange from LUF5843 to ZM241385. | X-RAY |
| 6PT2 | Crystal structure of the active delta opioid receptor in complex with the peptide agonist KGCHM07 | X-RAY |
| 6RZ6 | Crystal structure of the human cysteinyl leukotriene receptor 2 in complex with ONO-2570366 (C2221 space group) | X-RAY |
| 6RZ7 | Crystal structure of the human cysteinyl leukotriene receptor 2 in complex with ONO-2570366 (F222 space group) | X-RAY |
| 6RZ9 | Crystal structure of the human cysteinyl leukotriene receptor 2 in complex with ONO-2770372 | X-RAY |
| 6PT0 | Cryo-EM structure of human cannabinoid receptor 2-Gi protein in complex with agonist WIN 55,212-2 | CYRO-EM |
| 6OS1 | Structure of synthetic nanobody-stabilized angiotensin II type 1 receptor bound to TRV023 | X-RAY |
| 6OS2 | Structure of synthetic nanobody-stabilized angiotensin II type 1 receptor bound to TRV026 | X-RAY |
| 6VI4 | Nanobody-Enabled Monitoring of Kappa Opioid Receptor States | X-RAY |
| 6LW5 | Crystal structure of the human formyl peptide receptor 2 in complex with WKYMVm | X-RAY |
| 6WIV | Structure of human GABA(B) receptor in an inactive state | CYRO-EM |
| 6S0Q | Structure of the A2A adenosine receptor determined at SwissFEL using native-SAD at 4.57 keV from 50,000 diffraction patterns | X-RAY |
| 6LFM | Cryo-EM structure of a class A GPCR | CYRO-EM |
| 6LFO | Cryo-EM structure of a class A GPCR monomer | CYRO-EM |
| 7CFM | Cryo-EM structure of the P395-bound GPBAR-Gs complex | CYRO-EM |
| 7CFN | Cryo-EM structure of the INT-777-bound GPBAR-Gs complex | CYRO-EM |
| 6WGT | Crystal structure of HTR2A with hallucinogenic agonist | X-RAY |
| 6WH4 | Crystal structure of HTR2A with inverse agonist | X-RAY |
| 6XBJ | Structure of human SMO-D384R complex with Gi | CYRO-EM |
| 6XBK | Structure of human SMO-G111C/I496C complex with Gi | CYRO-EM |
| 6XBL | Structure of human SMO-Gi complex with SAG | CYRO-EM |
| 6WQA | 2.0A angstrom A2a adenosine receptor structure using XFEL data collected in helium atmosphere. | X-RAY |
| 7CZ5 | Cryo-EM structure of the human growth hormone-releasing hormone receptor-Gs protein complex | CYRO-EM |
| 6LPJ | A2AR crystallized in EROCOC17+4, LCP-SFX at 277 K | X-RAY |
| 6LPK | A2AR crystallized in EROCOC17+4, LCP-SFX at 293 K | X-RAY |
| 6LPL | A2AR crystallized in EROCOC17+4, SS-ROX at 100 K | X-RAY |
| 7BTS | Structure of human beta1 adrenergic receptor bound to epinephrine and nanobody 6B9 | X-RAY |
| 7BU6 | Structure of human beta1 adrenergic receptor bound to norepinephrine and nanobody 6B9 | X-RAY |
| 7BU7 | Structure of human beta1 adrenergic receptor bound to BI-167107 and nanobody 6B9 | X-RAY |
| 7BVQ | Structure of human beta1 adrenergic receptor bound to carazolol | X-RAY |
| 7D76 | Cryo-EM structure of the beclomethasone-bound adhesion receptor GPR97-Go complex | CYRO-EM |
| 7D77 | Cryo-EM structure of the cortisol-bound adhesion receptor GPR97-Go complex | CYRO-EM |
| 7ARO | Crystal structure of the non-ribose partial agonist LUF5833 bound to the adenosine A2A receptor | X-RAY |
| 7E14 | Compound2_GLP-1R_OWL833_Gs complex structure | CYRO-EM |
| 7DW9 | Cryo-EM structure of human V2 vasopressin receptor in complex with an Gs protein | CYRO-EM |
| 7F9Y | ghrelin-bound ghrelin receptor in complex with Gq | CYRO-EM |
| 7RM5 | MicroED structure of the human adenosine receptor at 2.8A | ELECTRON CRYSTALLOGRAPHY |
| 7V3Z | Structure of cannabinoid receptor type 1(CB1) | X-RAY |
| 7VDH | Cryo-EM structure of pseudoallergen receptor MRGPRX2 complex with C48/80, state2 | CYRO-EM |
| 7VDL | Cryo-EM structure of pseudoallergen receptor MRGPRX2 complex with circular cortistatin-14 | CYRO-EM |
| 7VDM | Cryo-EM structure of pseudoallergen receptor MRGPRX2 complex with substance P | CYRO-EM |
| 7VUY | Cryo-EM structure of pseudoallergen receptor MRGPRX2 complex with PAMP-12. state1 | CYRO-EM |
| 7VUZ | Cryo-EM structure of pseudoallergen receptor MRGPRX2 complex with PAMP-12, state2 | CYRO-EM |
| 7VV0 | Cryo-EM structure of pseudoallergen receptor MRGPRX2 complex with PAMP-12, local | CYRO-EM |
| 7VV3 | Cryo-EM structure of pseudoallergen receptor MRGPRX2 complex with linear cortistatin-14 | CYRO-EM |
| 7VV4 | Cryo-EM structure of pseudoallergen receptor MRGPRX2 complex with linear cortistatin-14, local | CYRO-EM |
| 7VV5 | Cryo-EM structure of pseudoallergen receptor MRGPRX2 complex with C48/80, state1 | CYRO-EM |
| 7VV6 | Cryo-EM structure of pseudoallergen receptor MRGPRX2 complex with C48/80 (local) | CYRO-EM |
| 7LI7 | apo serotonin transporter reconstituted in lipid nanodisc in presence of NaCl in occluded conformation | CYRO-EM |
| 7LIA | 5-HT bound serotonin transporter reconstituted in lipid nanodisc in presence of NaCl in outward facing conformation | CYRO-EM |
| 7MGW | 5-HT bound serotonin transporter reconstituted in lipid nanodisc in NaCl in occluded conformation | CYRO-EM |
| 7NA7 | Structures of human ghrelin receptor-Gi complexes with ghrelin and a synthetic agonist | CYRO-EM |
| 7NA8 | Structures of human ghrelin receptor-Gi complexes with ghrelin and a synthetic agonist | CYRO-EM |
| 7P00 | Human Neurokinin 1 receptor (NK1R) substance P Gq chimera (mGsqi) complex | CYRO-EM |
| 7P02 | Human Neurokinin 1 receptor (NK1R) substance P Gs complex | CYRO-EM |
| 7VOD | Crystal structure of 5-HT2AR in complex with cariprazine | X-RAY |
| 7VOE | Crystal structure of 5-HT2AR in complex with aripiprazole | X-RAY |
| 7F6H | Cryo-EM structure of human bradykinin receptor BK2R in complex Gq proteins and bradykinin | CYRO-EM |
| 7F6I | Cryo-EM structure of human bradykinin receptor BK2R in complex Gq proteins and kallidin | CYRO-EM |
| 7RKM | Structure of CX3CL1-US28-Gi-scFv16 in C-state | CYRO-EM |
| 7WC4 | Crystal structure of serotonin 2A receptor in complex with serotonin | X-RAY |
| 7WC5 | Crystal structure of serotonin 2A receptor in complex with psilocin | X-RAY |
| 7WC6 | Crystal structure of serotonin 2A receptor in complex with LSD | X-RAY |
| 7WC7 | Crystal structure of serotonin 2A receptor in complex with lisuride | X-RAY |
| 7WC8 | Crystal structure of serotonin 2A receptor in complex with lumateperone | X-RAY |
| 7WC9 | Crystal structure of serotonin 2A receptor in complex with non-hallucinogenic psychedelic analog | X-RAY |
| 7PX4 | Crystal structure of the adenosine A2A receptor (A2A-PSB1-bRIL) in complex with preladenant conjugate PSB-2113 | X-RAY |
| 7PYR | Crystal structure of the adenosine A2A receptor (A2A-PSB1-bRIL) in complex with preladenant conjugate PSB-2115 | X-RAY |
| 7VGY | Melatonin receptor1-2-Iodomelatonin-Gicomplex | CYRO-EM |
| 7VGZ | MT1-remalteon-Gi complex | CYRO-EM |
| 7VL8 | Cryo-EM structure of the Apo CCR1-Gi complex | CYRO-EM |
| 7VL9 | Cryo-EM structure of the CCL15(26-92) bound CCR1-Gi complex | CYRO-EM |
| 7VLA | Cryo-EM structure of the CCL15(27-92) bound CCR1-Gi complex | CYRO-EM |
| 7EJX | Structure of the GPR88-Gi1 signaling complex bound to a synthetic ligand | CYRO-EM |
| 7WQ3 | Galanin-bound galanin receptor 1 in complex with Gi | CYRO-EM |
| 7WQ4 | Galanin-bound galanin receptor 2 in complex with Gq | CYRO-EM |
| 7WU4 | Cryo-EM structure of the adhesion GPCR ADGRF1 in complex with miniGi | CYRO-EM |
| 7WU5 | Cryo-EM structure of the adhesion GPCR ADGRF1(H565A/T567A) in complex with miniGi | CYRO-EM |
| 7FEE | Crystal Structure of protein l | X-RAY |
| 7X2D | Cryo-EM structure of the tavapadon-bound D1 dopamine receptor and mini-Gs complex | CYRO-EM |
| 7X2F | Cryo-EM structure of the dopamine and LY3154207-bound D1 dopamine receptor and mini-Gs complex | CYRO-EM |
| 7ZI0 | Structure of human Smoothened in complex with cholesterol and SAG | X-RAY |
| 7XT8 | Serotonin 4 (5-HT4) receptor-Gs-Nb35 complex | CYRO-EM |
| 7XT9 | Serotonin 4 (5-HT4) receptor-Gs complex | CYRO-EM |
| 7XTC | Serotonin 7 (5-HT7) receptor-Gs-Nb35 complex | CYRO-EM |
| 8CXO | Cryo-EM structure of the unliganded mSMO-PGS2 in a lipidic environment | CYRO-EM |
| 8DU3 | Crystal structure of A2AAR-StaR2-bRIL in complex with compound 21a | X-RAY |
| 7XW6 | TSHR-Gs-M22 antibody-ML109 complex | CYRO-EM |
| 7XW5 | TSHR-thyroid stimulating hormone-Gs-ML109 complex | CYRO-EM |
| 8DPF | Cryo-EM structure of the 5HT2C receptor (INI isoform) bound to lorcaserin | CYRO-EM |
| 8DPG | Cryo-EM structure of the 5HT2C receptor (INI isoform) bound to psilocin | CYRO-EM |
| 8DPH | Cryo-EM structure of the 5HT2C receptor (VGV isoform) bound to lorcaserin | CYRO-EM |
| 8DPI | Cryo-EM structure of the 5HT2C receptor (VSV isoform) bound to lorcaserin | CYRO-EM |
| 8CU6 | Crystal structure of A2AAR-StaR2-S277-bRIL in complex with a novel A2a antagonist, LJ-4517 | X-RAY |
| 8CU7 | Crystal structure of A2AAR-StaR2-bRIL in complex with a novel A2a antagonist, LJ-4517 | X-RAY |
| 7F61 | Crystal structure of human histamine receptor H3R in complex with antagonist PF03654746 | X-RAY |
| 7PP1 | Crystal structure of the P2Y12 receptor in complex with the inverse agonist selatogrel. | X-RAY |
| 7T2G | CryoEM structure of mu-opioid receptor - Gi protein complex bound to mitragynine pseudoindoxyl (MP) | CYRO-EM |
| 8HDO | Structure of A2BR bound to synthetic agonists BAY 60-6583 | CYRO-EM |
| 8HDP | Structure of A2BR bound to endogenous agonists adenosine | CYRO-EM |
| 8H8J | Lodoxamide-bound GPR35 in complex with G13 | CYRO-EM |
| 8FYN | MicroED structure of A2A from plasma milled lamellae | ELECTRON CRYSTALLOGRAPHY |
| 8GNE | Crystal structure of human adenosine A2A receptor in complex with an insurmountable inverse agonist, KW-6356. | X-RAY |
| 7F6G | Cryo-EM structure of human angiotensin receptor AT1R in complex Gq proteins and Sar1-AngII | CYRO-EM |
| 8HK2 | C3aR-Gi-C3a protein complex | CYRO-EM |
| 8IRV | Dopamine Receptor D5R-Gs-Rotigotine complex | CYRO-EM |
| 8IRR | Dopamine Receptor D1R-Gs-Rotigotine complex | CYRO-EM |
| 8IRU | Dopamine Receptor D4R-Gi-Rotigotine complex | CYRO-EM |
| 8JCW | Cryo-EM structure of mGlu2-mGlu3 heterodimer in presence of LY341495 and NAM563 (dimerization mode I) | CYRO-EM |
| 8JCY | Cryo-EM structure of mGlu2-mGlu3 heterodimer in presence of LY341495, NAM563, and LY2389575 (dimerization mode I) | CYRO-EM |
| 8JD0 | Cryo-EM structure of mGlu2-mGlu3 heterodimer in presence of NAM563 | CYRO-EM |
| 8JD1 | Cryo-EM structure of mGlu2-mGlu3 heterodimer in Rco state | CYRO-EM |
| 8JD3 | Cryo-EM structure of Gi1-bound mGlu2-mGlu3 heterodimer | CYRO-EM |
| 8JD5 | Cryo-EM structure of Gi1-bound mGlu2-mGlu4 heterodimer | CYRO-EM |
| 8C9W | Crystal structure of the adenosine A2A receptor (construct A2A-PSB2-bRIL) complexed with Etrumadenant at the orthosteric pocket | X-RAY |
| 8H0Q | Structure of the GRP14-27-GRPR-Gq complex | CYRO-EM |
| 8A2O | Room-temperature structure of the stabilised A2A-Theophylline complex determined by synchrotron serial crystallography | X-RAY |
| 8A2P | Room-temperature structure of the stabilised A2A-LUAA47070 complex determined by synchrotron serial crystallography | X-RAY |
| 8JHY | Cryo-EM structure of compound 9n bound ketone body receptor HCAR2-Gi signaling complex | CYRO-EM |
| 8JII | Cryo-EM structure of compound 9n and niacin bound ketone body receptor HCAR2-Gi signaling complex | CYRO-EM |
| 8JIL | Cryo-EM structure of niacin bound ketone body receptor HCAR2-Gi signaling complex | CYRO-EM |
| 8JIM | Cryo-EM structure of MMF bound ketone body receptor HCAR2-Gi signaling complex | CYRO-EM |
| 8KH4 | Cryo-EM structure of the GPR161-Gs complex | CYRO-EM |
| 8KH5 | Cryo-EM structure of the GPR174-Gs complex bound to endogenous lysoPS | CYRO-EM |
| 7YFC | Cryo-EM structure of the histamine-bound histamine H4 receptor and Gq complex | CYRO-EM |
| 7YFD | Cryo-EM structure of the imetit-bound histamine H4 receptor and Gq complex | CYRO-EM |
| 8PWN | Structure of A2A adenosine receptor A2AR-StaR2-bRIL, solved at wavelength 2.75 A | X-RAY |
| 8G05 | Cryo-EM structure of an orphan GPCR-Gi protein signaling complex | CYRO-EM |
| 8W88 | Cryo-EM structure of the SEP363856-bound TAAR1-Gs complex | CYRO-EM |
| 8W8B | Cryo-EM structure of SEP-363856 bounded serotonin 1A (5-HT1A) receptor-Gi protein complex | CYRO-EM |
| 8WPU | Human calcium-sensing receptor(CaSR) bound to cinacalcet in complex with Gq protein | CYRO-EM |
| 8J6P | Cryo-EM structure of the MK-6892-bound human HCAR2-Gi1 complex | CYRO-EM |
| 8J6Q | Cryo-EM structure of the 3-HB and compound 9n-bound human HCAR2-Gi1 complex | CYRO-EM |
| 8T3S | Cryo-EM structure of the Butyrate bound FFA2-Gq complex | CYRO-EM |
| 8T3V | Cryo-EM structure of the DHA bound FFA1-Gq complex | CYRO-EM |
| 8SMV | GPR161 Gs heterotrimer | CYRO-EM |
| 8JT6 | 5-HT1A-Gi in complex with compound (R)-IHCH-7179 | CYRO-EM |
| 8JT8 | Crystal structure of 5-HT2AR in complex with (R)-IHCH-7179 | X-RAY |

```
sp|P41586-5|PACR_HUMAN      MAGVVHVSLAALLLLPMAPAMHSDCIFKKEQAMCLEKIQRANELMGFNDSSP--------	52
sp|P41586-3|PACR_HUMAN      MAGVVHVSLAALLLLPMAPAMHSDCIFKKEQAMCLEKIQRANELMGFNDSSPGCPGMWDN	60
sp|P41586|PACR_HUMAN        MAGVVHVSLAALLLLPMAPAMHSDCIFKKEQAMCLEKIQRANELMGFNDSSPGCPGMWDN	60
sp|P41586-2|PACR_HUMAN      MAGVVHVSLAALLLLPMAPAMHSDCIFKKEQAMCLEKIQRANELMGFNDSSPGCPGMWDN	60
sp|P41586-4|PACR_HUMAN      MAGVVHVSLAALLLLPMAPAMHSDCIFKKEQAMCLEKIQRANELMGFNDSSPGCPGMWDN	60
                            ****************************************************

sp|P41586-5|PACR_HUMAN      -------------------------------------------------DMGVVSRNCTE	63
sp|P41586-3|PACR_HUMAN      ITCWKPAHVGEMVLVSCPELFRIFNPDQ---------------------DMGVVSRNCTE	99
sp|P41586|PACR_HUMAN        ITCWKPAHVGEMVLVSCPELFRIFNPDQVWETETIGESDFGDSNSLDLSDMGVVSRNCTE	120
sp|P41586-2|PACR_HUMAN      ITCWKPAHVGEMVLVSCPELFRIFNPDQVWETETIGESDFGDSNSLDLSDMGVVSRNCTE	120
sp|P41586-4|PACR_HUMAN      ITCWKPAHVGEMVLVSCPELFRIFNPDQ---------------------DMGVVSRNCTE	99
                                                                             ***********

sp|P41586-5|PACR_HUMAN      DGWSEPFPHYFDACGFDEYESETGDQDYYYLSVKALYTVGYSTSLVTLTTAMVILCRFRK	123
sp|P41586-3|PACR_HUMAN      DGWSEPFPHYFDACGFDEYESETGDQDYYYLSVKALYTVGYSTSLVTLTTAMVILCRFRK	159
sp|P41586|PACR_HUMAN        DGWSEPFPHYFDACGFDEYESETGDQDYYYLSVKALYTVGYSTSLVTLTTAMVILCRFRK	180
sp|P41586-2|PACR_HUMAN      DGWSEPFPHYFDACGFDEYESETGDQDYYYLSVKALYTVGYSTSLVTLTTAMVILCRFRK	180
sp|P41586-4|PACR_HUMAN      DGWSEPFPHYFDACGFDEYESETGDQDYYYLSVKALYTVGYSTSLVTLTTAMVILCRFRK	159
                            ************************************************************

sp|P41586-5|PACR_HUMAN      LHCTRNFIHMNLFVSFMLRAISVFIKDWILYAEQDSNHCFISTVECKAVMVFFHYCVVSN	183
sp|P41586-3|PACR_HUMAN      LHCTRNFIHMNLFVSFMLRAISVFIKDWILYAEQDSNHCFISTVECKAVMVFFHYCVVSN	219
sp|P41586|PACR_HUMAN        LHCTRNFIHMNLFVSFMLRAISVFIKDWILYAEQDSNHCFISTVECKAVMVFFHYCVVSN	240
sp|P41586-2|PACR_HUMAN      LHCTRNFIHMNLFVSFMLRAISVFIKDWILYAEQDSNHCFISTVECKAVMVFFHYCVVSN	240
sp|P41586-4|PACR_HUMAN      LHCTRNFIHMNLFVSFMLRAISVFIKDWILYAEQDSNHCFISTVECKAVMVFFHYCVVSN	219
                            ************************************************************

sp|P41586-5|PACR_HUMAN      YFWLFIEGLYLFTLLVETFFPERRYFYWYTIIGWGTPTVCVTVWATLRLYFDDTGCWDMN	243
sp|P41586-3|PACR_HUMAN      YFWLFIEGLYLFTLLVETFFPERRYFYWYTIIGWGTPTVCVTVWATLRLYFDDTGCWDMN	279
sp|P41586|PACR_HUMAN        YFWLFIEGLYLFTLLVETFFPERRYFYWYTIIGWGTPTVCVTVWATLRLYFDDTGCWDMN	300
sp|P41586-2|PACR_HUMAN      YFWLFIEGLYLFTLLVETFFPERRYFYWYTIIGWGTPTVCVTVWATLRLYFDDTGCWDMN	300
sp|P41586-4|PACR_HUMAN      YFWLFIEGLYLFTLLVETFFPERRYFYWYTIIGWGTPTVCVTVWATLRLYFDDTGCWDMN	279
                            ************************************************************

sp|P41586-5|PACR_HUMAN      DSTALWWVIKGPVVGSIMVNFVLFIGIIVILVQKLQSPDMGGNESSIY------------	291
sp|P41586-3|PACR_HUMAN      DSTALWWVIKGPVVGSIMVNFVLFIGIIVILVQKLQSPDMGGNESSIY------------	327
sp|P41586|PACR_HUMAN        DSTALWWVIKGPVVGSIMVNFVLFIGIIVILVQKLQSPDMGGNESSIY------------	348
sp|P41586-2|PACR_HUMAN      DSTALWWVIKGPVVGSIMVNFVLFIGIIVILVQKLQSPDMGGNESSIYFSCVQKCYCKPQ	360
sp|P41586-4|PACR_HUMAN      DSTALWWVIKGPVVGSIMVNFVLFIGIIVILVQKLQSPDMGGNESSIYFSCVQKCYCKPQ	339
                            ************************************************

sp|P41586-5|PACR_HUMAN      ----------------LRLARSTLLLIPLFGIHYTVFAFSPENVSKRERLVFELGLGSFQ	335
sp|P41586-3|PACR_HUMAN      ----------------LRLARSTLLLIPLFGIHYTVFAFSPENVSKRERLVFELGLGSFQ	371
sp|P41586|PACR_HUMAN        ----------------LRLARSTLLLIPLFGIHYTVFAFSPENVSKRERLVFELGLGSFQ	392
sp|P41586-2|PACR_HUMAN      RAQQHSCKMSELSTITLRLARSTLLLIPLFGIHYTVFAFSPENVSKRERLVFELGLGSFQ	420
sp|P41586-4|PACR_HUMAN      RAQQHSCKMSELSTITLRLARSTLLLIPLFGIHYTVFAFSPENVSKRERLVFELGLGSFQ	399
                                            ********************************************

sp|P41586-5|PACR_HUMAN      GFVVAVLYCFLNGEVQAEIKRKWRSWKVNRYFAVDFKHRHPSLASSGVNGGTQLSILSKS	395
sp|P41586-3|PACR_HUMAN      GFVVAVLYCFLNGEVQAEIKRKWRSWKVNRYFAVDFKHRHPSLASSGVNGGTQLSILSKS	431
sp|P41586|PACR_HUMAN        GFVVAVLYCFLNGEVQAEIKRKWRSWKVNRYFAVDFKHRHPSLASSGVNGGTQLSILSKS	452
sp|P41586-2|PACR_HUMAN      GFVVAVLYCFLNGEVQAEIKRKWRSWKVNRYFAVDFKHRHPSLASSGVNGGTQLSILSKS	480
sp|P41586-4|PACR_HUMAN      GFVVAVLYCFLNGEVQAEIKRKWRSWKVNRYFAVDFKHRHPSLASSGVNGGTQLSILSKS	459
                            ************************************************************

sp|P41586-5|PACR_HUMAN      SSQIRMSGLPADNLAT	411
sp|P41586-3|PACR_HUMAN      SSQIRMSGLPADNLAT	447
sp|P41586|PACR_HUMAN        SSQIRMSGLPADNLAT	468
sp|P41586-2|PACR_HUMAN      SSQIRMSGLPADNLAT	496
sp|P41586-4|PACR_HUMAN      SSQIRMSGLPADNLAT	475
```

**Figure S1: Sequence Alignment of PAC1R Variants. PAC1null (P41586) and PAC1hop (P41586-2) differ in the ICL3 region.**

**Figure S2: All-atom Contact Maps (left) vs. CG Contact Maps (right).** An active and inactive model of the PAC1R (null) with GM3 contacts within 8Å color coded. MARTINI contact heatmaps are ranked from white to red, while all-atom REMD contact heatmaps are ranked from purple to red. ECD contacts are observed to be heavily favored in both inactive models.
